# *In situ* architecture of neuronal α-Synuclein inclusions

**DOI:** 10.1101/2020.08.07.234138

**Authors:** Victoria A. Trinkaus, Irene Riera-Tur, Antonio Martínez-Sánchez, Felix J.B. Bäuerlein, Qiang Guo, Thomas Arzberger, Wolfgang Baumeister, Irina Dudanova, Mark S. Hipp, F. Ulrich Hartl, Rubén Fernández-Busnadiego

## Abstract

α-Synuclein (α-Syn) aggregation is a hallmark of devastating neurodegenerative disorders including Parkinson’s disease (PD) and multiple systems atrophy (MSA)^1,2^. α-Syn aggregates spread throughout the brain during disease progression^2^, suggesting mechanisms of intercellular seeding. Formation of α-Syn amyloid fibrils is observed *in vitro*^3,4^ and fibrillar α-Syn has been purified from patient brains^5,6^, but recent reports questioned whether disease-relevant α-Syn aggregates are fibrillar in structure^7-9^. Here we use cryo-electron tomography (cryo-ET) to image neuronal Lewy body-like α-Syn inclusions *in situ* at molecular resolution. We show that the inclusions consist of α-Syn fibrils crisscrossing a variety of cellular organelles such as the endoplasmic reticulum (ER), mitochondria and autophagic structures, without interacting with membranes directly. Neuronal inclusions seeded by recombinant or MSA patient-derived α-Syn aggregates have overall similar architecture, although MSA-seeded fibrils show higher structural flexibility. Using gold-labeled seeds we find that aggregate nucleation is predominantly mediated by α-Syn oligomers, with fibrils growing unidirectionally from the seed. Our results conclusively demonstrate that neuronal α-Syn inclusions contain α-Syn fibrils intermixed with cellular membranes, and illuminate the mechanism of aggregate nucleation.

## Main

Early electron microscopy (EM) studies suggested that the Lewy body inclusions characteristic of PD are fibrillar^10,11^. However, conventional EM lacks the resolution to unequivocally determine the molecular identity of Lewy body fibrils *in situ*, a problem further complicated by the cross-reactivity of α-Syn antibodies with neurofilaments^12^. A recent study using correlative EM on chemically fixed PD brain tissue suggested that cellular membranes were the main component of Lewy bodies, alongside with unidentified fibrillar material^7^. These findings resonated with reports^13^ that native α-Syn binds lipids such as synaptic vesicle membranes^14^, observations that lipids can catalyze α-Syn aggregation *in vitro*^15^, and that α-Syn expression in cells is associated with membrane abnormalities^9^. Thus, the disease relevance of fibrillar (amyloid-like) α-Syn aggregation has been questioned, leading to a model in which the main role of α-Syn in Lewy bodies is to cluster cellular membranes^8,9^. Cryo-ET is ideally positioned to test these new ideas, as it can reveal the molecular architecture of protein aggregates at high resolution within neurons pristinely preserved by vitrification^16-18^.

We performed cryo-ET on neuronal α-Syn aggregates using a well-established seeding paradigm that recapitulates key features of pathological Lewy bodies and their inter-neuronal spreading^19^. Primary mouse neurons were cultured on EM grids, transduced with GFP-α-Syn and incubated with recombinant α-Syn pre-formed fibrils (PFFs) (Extended Data Fig. 1a). Unless otherwise stated, all experiments were carried out using the familial A53T α-Syn mutation due to its higher seeding potency^20^. As reported^19^, seeding of neurons led to the formation of GFP-α-Syn inclusions that were positive for Lewy body markers including phospho-α-Syn (Ser129) and p62 (Extended Data Fig. 1b, c). GFP-α-Syn inclusions in cell bodies or neurites were targeted for cryo-ET by correlative microscopy and cryo-focused ion beam (cryo-FIB) milling^16-18,21,22^ (Extended Data Fig. 2). In all cases, this analysis revealed large fibrillar accumulations at sites of GFP-α-Syn fluorescence (Fig. 1a, d). Interestingly, the fibrils appeared to be composed of a core decorated by globular GFP-like densities (Fig. 1b), reminiscent of GFP-labeled polyQ and *C9orf72* poly-GA aggregates^16,17^, and were clearly distinct from cytoskeletal elements (Fig. 1c). Notably, these fibrillar accumulations were populated by numerous cellular organelles, including ER, mitochondria, autophagolysosomal structures and small vesicles (Fig. 1a, d). Thus, the α-Syn inclusions formed in our cellular system recapitulated the key ultrastructural features of Lewy bodies, consistent with recent reports^7,23^.

**Fig. 1.**
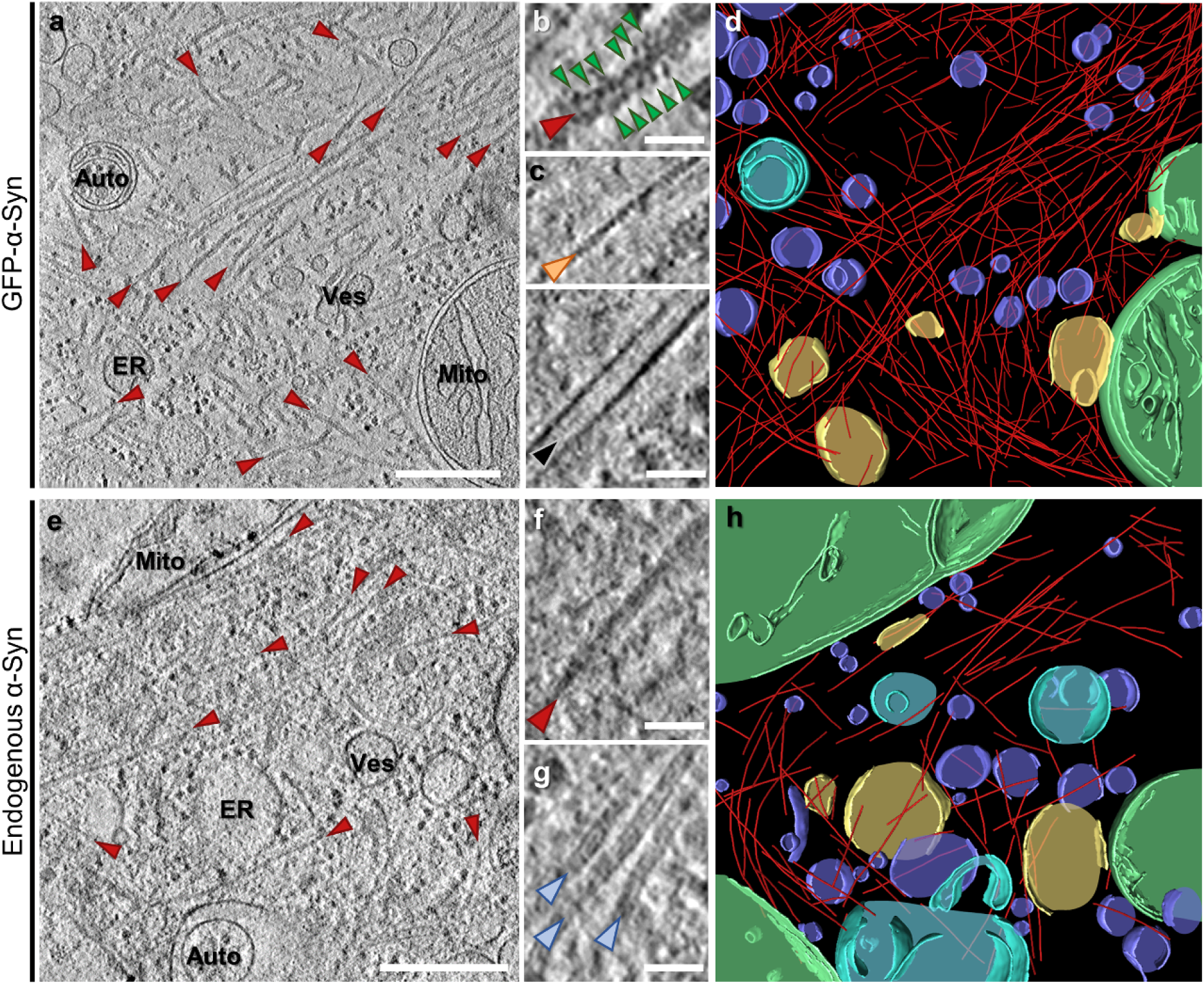
Cryo-ET imaging of α-Syn aggregates seeded by PFFs in neurons. **a**, A tomographic slice (thickness 1.8 nm) of an inclusion seeded by PFFs in a neuron expressing GFP-α-Syn. Auto: autophagosome; ER: endoplasmic reticulum; Mito: mitochondrion; Ves: vesicles. Fibrils are marked by red arrowheads. Scale bar: 350 nm. **b**, Magnified view of a fibril with GFP-like densities (green arrowheads) decorating the fibril core. Scale bar: 30 nm. **c**, Magnified views of an actin filament (orange arrowhead) and a microtubule (black arrowhead). Scale bar: 30 nm. **d**, 3D rendering of **a** showing α-Syn fibrils (red), an autophagosome (cyan), ER (yellow), mitochondria (green) and various vesicles (purple). **e**, A tomographic slice (thickness 1.4 nm) of an inclusion seeded by PFFs in a neuron expressing p62-RFP. Scale bar: 350 nm. **f**, Magnified view of a fibril. Note that fibrils in cells not expressing GFP-α-Syn are not decorated by GFP-like densities. Scale bar: 30 nm. **g**, Magnified view of neurofilaments (blue arrowheads). Scale bar: 30 nm. **h**, 3D rendering of **e**.

**Fig. 2.**
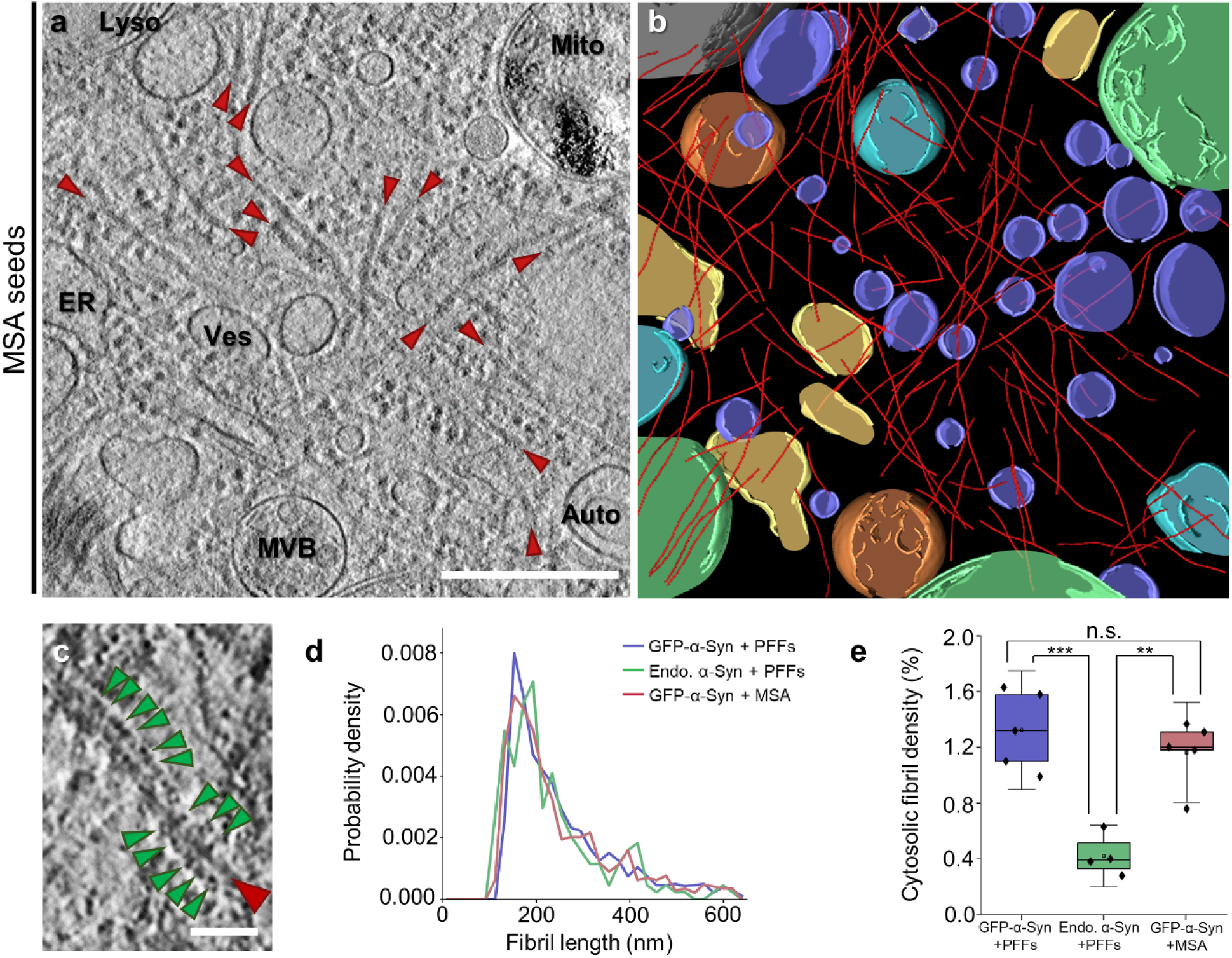
Cryo-ET imaging of α-Syn aggregates seeded by MSA patient brain material in neurons. **a**, A tomographic slice (thickness 1.4 nm) of an inclusion seeded by MSA patient brain material in a neuron expressing GFP-α-Syn. Auto: autophagosome; ER: endoplasmic reticulum; Lyso: lysosome; Mito: mitochondrion; MVB: multivesicular body; Ves: vesicles. Fibrils are marked by red arrowheads. Scale bar: 350 nm. **b**, 3D rendering of **a** showing α-Syn fibrils (red), autophagosomes (cyan), ER (yellow), a lysosome (grey), mitochondria (green), multivesicular bodies (orange) and various vesicles (purple). **c**, Magnified view of a fibril with GFP-like densities (green arrowheads) decorating the fibril core. Scale bar: 30 nm. **d**, Histogram of fibril length. N = 1471 (GFP-α-Syn + PFFs), 220 (endogenous α-Syn + PFFs) and 721 (GFP-α-Syn + MSA) fibrils analyzed. See also Extended Data Table 1. **e**, Box plots of cytosolic fibril density within inclusions. The horizontal lines of each box represent 75% (top), 50% (middle) and 25% (bottom) of the values, and a black square the average value. Whiskers represent standard deviation and black diamonds the individual data points. N = 5 (GFP-α-Syn + PFFs), 4 (endogenous α-Syn + PFFs) and 5 (GFP-α-Syn + MSA) tomograms analyzed; n.s., ** and *** indicate respectively p = 0.4, p = 0.0010 and p = 7*10^−4^ by one-way ANOVA. See also Extended Data Table 1.

To further investigate the nature of the fibrils observed at sites of GFP-α-Syn fluorescence and avoid possible artifacts caused by GFP-α-Syn overexpression, we next imaged inclusions formed by endogenous α-Syn in neurons seeded by recombinant PFFs. Given the high p62 signal observed in Lewy bodies^1,24^ and GFP-α-Syn inclusions (Extended Data Fig. 1c), we expressed p62-RFP as a surrogate marker^17^ of endogenous α-Syn inclusions (Extended Data Fig. 1d) to guide correlative cryo-FIB/ET analysis. Although endogenous α-Syn inclusions were smaller than those formed by GFP-α-Syn (Extended Data Fig. 1b), cryo-ET imaging revealed a similar nanoscale organization, consisting of cellular membranes crisscrossed by abundant fibrils (Fig. 1e, h). Importantly, the fibrils appeared identical to those observed in GFP-α-Syn inclusions (Fig. 1b), except that they were not decorated by globular densities (Fig. 1f). The fibrils were also clearly distinct from neurofilaments (Fig. 1g). These data conclusively demonstrate that the fibrils observed in α-Syn inclusions are formed by α-Syn, and argue against a major effect of GFP-α-Syn overexpression on inclusion architecture. Nevertheless, GFP-α-Syn overexpression enhanced the rate of inclusion formation and neuronal toxicity (Extended Data Fig. 1e, f), implicating α-Syn aggregates in neuronal death.

Recent studies have demonstrated that amyloid fibrils, including those formed by α-Syn, may adopt different conformations when purified from patient brain in comparison to fibrils generated *in vitro* from recombinant proteins^25,26^. Therefore, to assess the disease relevance of our findings using recombinant PFFs, we seeded primary neurons expressing GFP-α-Syn with α-Syn aggregates purified from MSA patient brain (Extended Data Fig. 3). Similar to PFFs, MSA seeds triggered the formation of intracellular GFP-α-Syn inclusions positive for phospho-α-Syn (Ser129) and p62 (Extended Data Fig. 3e). Most importantly, cryo-ET analysis showed that MSA-seeded neuronal aggregates also consisted of a dense meshwork of α-Syn fibrils interspersed by cellular organelles (Fig. 2a, b, c). Therefore, our results show that neuronal α-Syn aggregates seeded by patient material are formed by accumulations of α-Syn fibrils and cellular membranes.

**Fig. 3.**
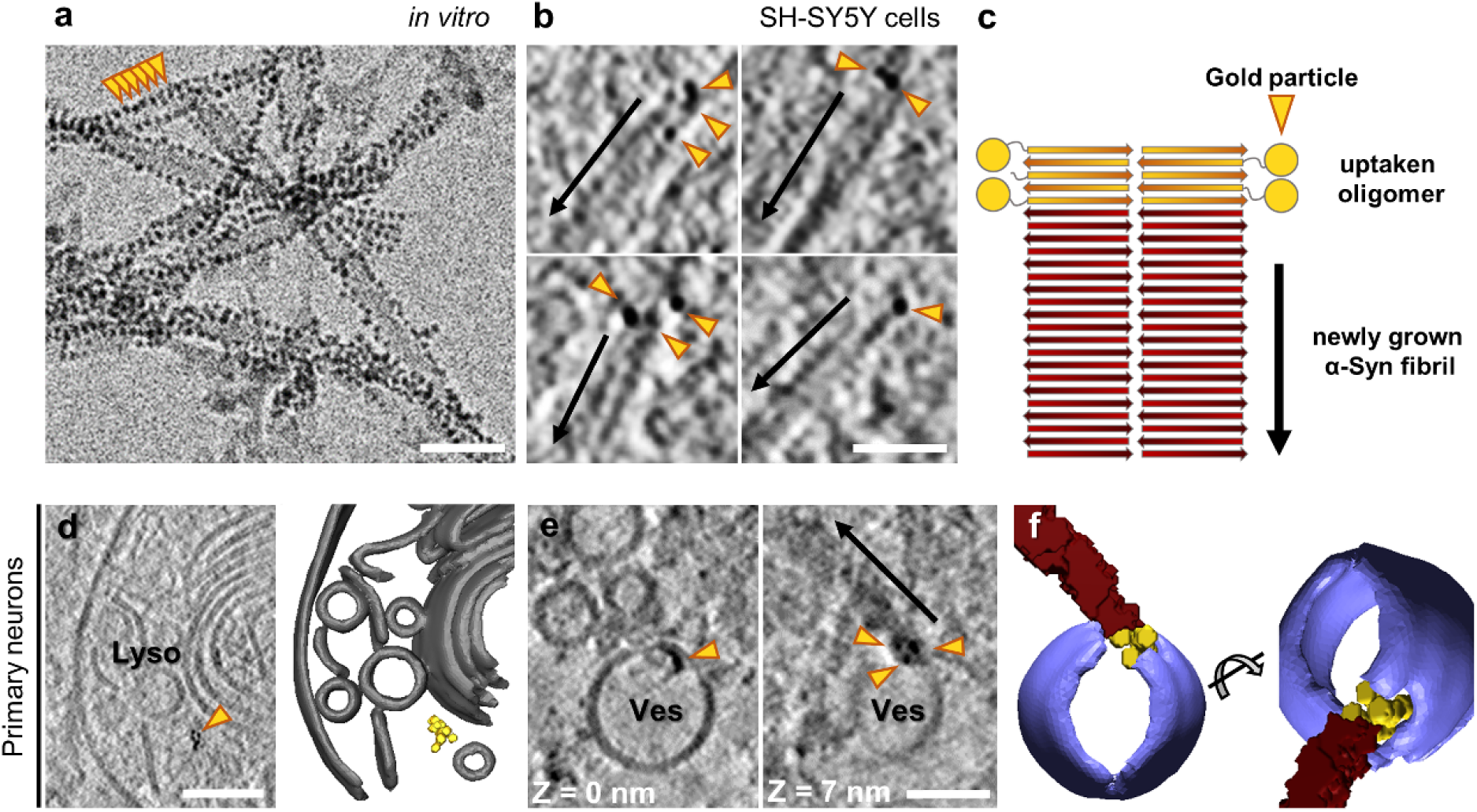
Seeding of α-Syn aggregates by gold-labeled PFFs. **a**, Cryo-electron microscopy image of PFFs labeled with 3 nm-gold beads (orange arrowheads) via NHS-esterification. Scale bar: 30 nm. **b**, Tomographic slices (thickness 1.8 nm) showing α-Syn fibrils nucleated by gold-labeled PFFs within SH-SY5Y cells expressing GFP-α-Syn. Arrows mark the direction of fibril growth from the gold-labeled seed. Scale bar: 40 nm. **c**, Schematic of the hypothetical molecular organization of α-Syn fibrils nucleated by gold-labeled PFFs. **d**, A tomographic slice (thickness 1.4 nm; left) and 3D rendering (right) showing gold-labeled PFFs within the lumen of a lysosome in a primary neuron expressing GFP-α-Syn. Lyso: lysosome. Lysosomal membranes (grey), gold particles labeling the PFF (yellow). Scale bar: 70 nm. **e**, Tomographic slices (thickness 1.4 nm) at different Z heights showing gold labeled PFFs found within the membrane of a vesicle (Ves) and nucleating an α-Syn fibril (arrow) in a primary neuron expressing GFP-α-Syn. Scale bar: 30 nm. **f**, 3D rendering of **e** in two different orientations. Vesicle membrane (purple), α-Syn fibril (red), gold particles (yellow).

We further investigated possible morphological differences between fibrils seeded by PFFs and MSA aggregates, and in neurons expressing endogenous α-Syn or GFP-α-Syn. In all cases mean fibril length was ∼250 nm (Fig. 2d, Extended Data Table 1). However, fibril density within inclusions was significantly higher in cells expressing GFP-α-Syn (Fig. 2e, Extended Data Table 1), likely due to the higher expression level of this construct, resulting in a higher aggregate load (Extended Data Fig. 1b, e). We next calculated the persistence length of the fibrils to investigate their mechanical properties. Interestingly, whereas PFF-seeded fibrils in neurons expressing GFP-α-Syn or endogenous α-Syn were almost identical in persistence length (Extended Data Fig. 4), MSA-seeded GFP-α-Syn fibrils displayed a considerably lower persistence length (Extended Data Fig. 4), reflecting higher structural flexibility. These values are in the range of those measured for α-Syn^27^ and tau^28^ fibrils *in vitro*, as well as for polyQ fibrils *in situ*^16^. Our measurements are also consistent with single-particle studies reporting a higher twist, indicative of higher flexibility^29^, for MSA-derived fibrils compared to recombinant fibrils^26,30^. Thus, different types of exogenous α-Syn aggregates seed neuronal inclusions with different mechano-physical properties.

**Fig. 4.**
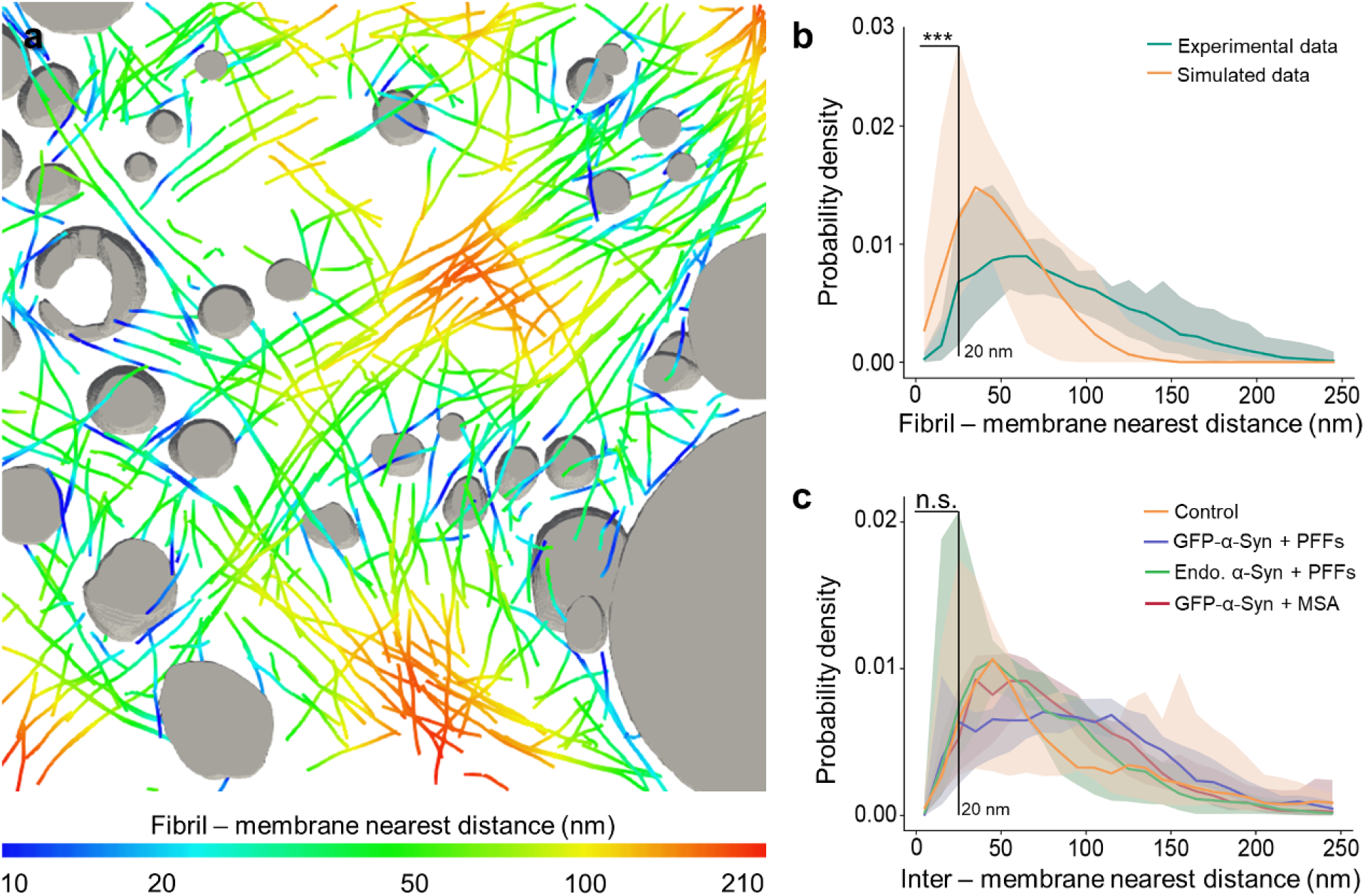
Quantification of fibril-membrane and inter-membrane distances within α-Syn inclusions. **a**, Visualization of fibril-membrane distances in the tomogram rendered in Fig. 1d. Organelles are shown in grey, and fibrils are color-coded according to their distance to the nearest organelle membrane. **b**, Histogram of nearest distances between a fibril and a membrane in the pooled experimental data (N = 14 tomograms, including 5 of GFP-α-Syn + PFFs, 4 of endogenous α-Syn + PFFs and 5 of GFP-α-Syn + MSA) and in simulations shifting and rotating fibrils from their experimentally determined positions (200 simulations for each experimental tomogram). Solid lines represent the median of all tomograms. The shaded areas represent 5-95% confidence intervals. Fibril-membrane distances < 20 nm are significantly more abundant in the simulated data (p = 2.4*10^−10^ by two-tailed Kolmogorov-Smirnov test). See also Extended Data Table 1. **c**, Histogram of inter-membrane nearest distances for all organellar membranes in the tomograms. Intermembrane distances are not significantly different within α-Syn inclusions than in control untransduced and unseeded cells (p = 0.754 by two-tailed Kolmogorov-Smirnov test). N = 5 (untransduced - PFFs), 5 (GFP-α-Syn + PFFs), 4 (endogenous α-Syn + PFFs) and 5 (GFP-α-Syn + MSA) tomograms analyzed. See also Extended Data Table 1.

The seeding of intracellular aggregation by extracellular aggregates may underlie the spreading of pathology across different brain regions during the progression of various neurodegenerative diseases, including synucleinopathies^31^. To gain a better mechanistic understanding of the seeding process, we tracked the fate of extracellular gold-labeled α-Syn seeds upon internalization into neurons expressing GFP-α-Syn. In this case, we used WT PFFs as they allowed higher labeling efficiency. Recombinant WT α-Syn fibrils were conjugated to 3-nm gold beads via NHS ester coupling, resulting in densely gold-labeled PFFs (Fig. 3a) that efficiently seeded the formation of neuronal GFP-α-Syn inclusions (Extended Data Fig. 5a). Some of these experiments were also carried out in a SH-SY5Y cell line stably expressing GFP-α-Syn as a simpler model system (Extended Data Fig. 6). Interestingly, cryo-ET analysis of inclusions seeded by gold-labeled PFFs showed GFP-α-Syn fibrils with one end decorated by 3-10 gold particles (Fig. 3b, c), indicating that exogenous seeds nucleate the fibrillation of cellular α-Syn in a polarized manner, consistent with the polarized structure of α-Syn fibrils^32^. These data also show that the nucleation-relevant seeds consist of oligomeric α-Syn. Therefore, despite the presence of abundant large fibrils in the exogenously added PFF material (Fig. 3a), these species are apparently not efficiently internalized. On the other hand, given the mechano-physical differences between neuronal fibrils growing from PFFs and MSA seeds (Extended Data Fig. 4), the seeding-competent oligomers likely contain the necessary information to confer these structural features. Gold-labeled α-Syn was also observed within the lumen of endolysosomal compartments (Fig. 3d, Extended Data Fig. 5b) and at their membrane (Fig. 3e, Extended Data Fig. 5b). Although the nucleation of α-Syn fibrils was occasionally observed directly at such membrane-bound gold-labeled structures (Fig. 3e, f), most gold-labeled fibrils were cytosolic (Fig. 3b). These data are in agreement with a model where oligomeric α-Syn seeds entering the cell are targeted to endosomes, from which they escape and trigger intracellular nucleation of α-Syn fibrils^33^.

The affinity of α-Syn for lipids^13^ has led to the proposal that α-Syn drives the accumulation of cellular membranes in Lewy bodies^7-9^, e.g. by fibril-membrane contacts as observed for polyQ fibrils^16^. Such contacts existed within α-Syn inclusions (Extended Data Fig. 7a, b), but they were extremely rare and did not seem to cause the kind of membrane deformations (Extended Data Fig. 7a) seen with polyQ^16^. Although we found a few examples where fibrils did contact membranes at areas of high curvature (Extended Data Fig. 7b), such areas also existed in the absence of fibril contacts (Extended Data Fig. 7c). Thus, apparent fibril-membrane contacts seemed to be mainly a consequence of the crowded cellular environment. To test this hypothesis, we computationally introduced random shifts and rotations to the experimentally determined positions of α-Syn fibrils within the tomograms. This analysis revealed that close fibril-membrane distances (< 20 nm) were significantly more frequent in random simulations than in the experimental data (Fig. 4a, b; Extended Data Fig. 7d, Extended Data Table 1). Together, these results indicate that direct interactions between α-Syn fibrils and membranes are infrequent and unlikely to induce substantial membrane clustering.

However, membrane clustering could also be driven by α-Syn species smaller than fibrils, which cannot be readily detected by cryo-ET. For example, soluble α-Syn molecules can cluster vesicles at distances shorter than 15 nm *in vitro*^34^. To explore this possibility, we compared the shortest distances between all cellular membranes in tomograms of α-Syn inclusions and in untransduced, unseeded control neurons. This analysis revealed that close contacts (<20 nm) between membranes were similarly common within α-Syn inclusions as in control cells (Fig. 4c; Extended Data Fig. 7e, Extended Data Table 1), arguing against α-Syn-mediated membrane clustering in inclusions.

Altogether, we show that neuronal α-Syn aggregates consist of both α-Syn fibrils and various cellular membranes, reconciling conflicting reports on the molecular architecture of Lewy bodies. Our findings strongly support the view that the unidentified fibrils observed in Lewy bodies of postmortem brain tissue^7^ are indeed α-Syn fibrils. Intracellular α-Syn aggregation can be triggered by internalized extracellular oligomeric seeds, suggesting that this mechanism underlies the spreading of aggregate pathology. However, α-Syn does not drive membrane clustering directly. Thus, the question why membrane structures are enriched in Lewy bodies remains to be addressed. An intriguing possibility is that vesicular organelles accumulate in Lewy bodies as a result of the impairment of the autophagic and endolysosomal machineries by α-Syn aggregation^35^.

## Methods

### Plasmids

Plasmids for the expression of recombinant α-Syn were: pT7-7 α-Syn (Addgene plasmid #36046^36^; http://n2t.net/addgene:36046; RRID:Addgene_36046) and pT7-7 α-Syn A53T (Addgene plasmid #105727^37^; http://n2t.net/addgene:105727; RRID:Addgene_105727) (gift from Hilal Lashuel).

Plasmid EGFP-α-SynA53T (Addgene plasmid #40823^38^; http://n2t.net/addgene:40823; RRID:Addgene_40823) was used for expression in SH-SY5Y cells (gift from David Rubinsztein).

The following plasmids were used for viral transfections: pFhSynW2^39^ (GFP-SynA53T-Flag, Flag-GFP), FU3a (p62-tagRFP)^17^, psPAX2 (a gift from Didier Trono; Addgene plasmid # 12260; http://n2t.net/addgene:12260; RRID:Addgene_12260) and pVsVg^40^. pFhSynW2 and pVsVg were a gift of Dieter Edbauer.

pFhSynW2 GFP-synA53T-Flag was cloned by inserting the GFP-α-SynA53T sequence from plasmid EGFP-α-SynA53T between the XmaI and NheI restriction sites using the following primers: forward: GCA GTC GAG AGG ATC CCG GGC CCA CCA TGG TGA GCA AGG GCG AG, and reverse: CCG CTC TAG AGC TAG CTT ATT TAT CGT CGT CAT CCT TGT AAT CGG CTT CAG GTT CGT AGT CTT GAT AC.

pFhSynW2 Flag-GFP was cloned by inserting the GFP sequence from the plasmid EGFP-α-SynA53T between the BamHI and EcoRI restriction sites using the following primers: forward: GAG CGC AGT CGA GAG GAT CCC CCA CCA TGG ATT ACA AGG ATG ACG ACG ATA AGC CCG GGA TGG TGA GCA AGG GCG AG, and reverse: GCT TGA TAT CGA ATT CTT ACT TGT ACA GCT CGT CCA TGC.

### Antibodies

The following primary antibodies were used: GFP (A10262, Thermo Fisher, 1:500; RRID: AB_2534023), K48-linked ubiquitin (05-1307, Millipore; 1:500; RRID: AB_1587578), MAP2 (NB300-213, Novus Biologicals; 1:500; RRID: AB_2138178), p62 (ab56416, Abcam; 1:200; RRID: AB_945626), phospho S129 α-Syn (ab51253, Abcam; 1:500 for immunofluorescence, 1:2500 for western blot; RRID: AB_869973), α-Syn (610787, BD Biosciences; 1:1000; RRID: AB_398108) and p62 lck ligand (610832, BD Biosciences; 1:100; RRID: AB_398151).

The following secondary antibodies were used: Alexa Fluor 488 AffiniPure Donkey Anti-Chicken (703-545-155, Jackson ImmunoResearch; 1:250), Alexa Fluor 647 AffiniPure Donkey Anti-Chicken (703-605-155, Jackson ImmunoResearch; 1:250), Cy3 AffiniPure Donkey Anti-Rabbit (711-165-152, Jackson ImmunoResearch; 1:250), Alexa Fluor 488 AffiniPure Donkey Anti-Mouse (715-545-150, Jackson ImmunoResearch; 1:250), Cy3-conjugated AffiniPure Goat anti-mouse IgG (115-165-003, Jackson ImmunoResearch; 1:1000), Cy3-conjugated AffiniPure Goat anti-rabbit (111-165-045, Dianova; 1:1000; RRID: AB_2338003), HRP-conjugated goat anti-rabbit (A9169, Sigma; 1:5000; RRID: AB_258434).

### Recombinant α-Syn purification and fibril assembly

Recombinant human WT and A53T α-Syn were purified similarly as described^19^. In brief, E. coli Rosetta-gami 2(DE3) cells (Novagen) were transformed with pT7-7 α-Syn or pT7-7 α-Syn A53T. Protein expression was induced by 1 mM IPTG for 4 h. Bacteria were harvested and pellets were lysed in high salt buffer (750 mM NaCl, 50 mM Tris, pH 7.6, 1 mM EDTA). The lysate was sonicated for 5 min and boiled subsequently. The boiled suspension was centrifuged, the supernatant dialyzed in 50 mM NaCl, 10 mM Tris and 1 mM EDTA and purified by size exclusion HPLC (Superdex 200). Fractions were collected and those containing α-Syn were combined. The combined fractions were applied onto an anion exchange column (MonoQ). α-Syn was purified by a gradient ranging from 50 mM to 1 M NaCl. Fractions containing α-Syn were combined and dialyzed in 150 mM KCl, 50 mM Tris, pH 7.6.

For fibril assembly, purified α-Syn monomers (5 mg/mL) were centrifuged at high speed (100,000 xg) for 1 h. The supernatant was transferred into a new reaction tube and incubated with constant agitation (1,000 rpm) at 37 °C for 24 h. The presence of α-Syn fibrils was confirmed by negative stain EM. Except for gold labeling experiments, cells were seeded using A53T α-Syn PFFs.

Labeling of fibrils with 3 nm monovalent gold-beads (Nanopartz) via NHS ester coupling was performed as described in the manufacturer’s protocol. In brief, WT α-Syn PFFs were dialyzed in PBS and subsequently added to the gold beads. The reaction was facilitated by constant agitation at 30 °C for 30 min. Labeled PFFs and free gold beads were separated by sequential centrifugation and washing with 0.1 % Tween20 and 1 % PBS. Labeling of PFFs with gold beads was confirmed by negative stain and cryo-EM.

### Immunohistochemistry on MSA patient brain

MSA patient brain tissue was obtained from Neurobiobank Munich (Germany). All autopsy cases of the Neurobiobank Munich are collected on the basis of an informed consent according to the guidelines of the ethics commission of the Ludwig-Maximilians-University Munich, Germany. The experiments performed in this paper were approved by the Max Planck Society’s Ethics Council. The sample was from a male patient who died at the age of 54, 6 years after being diagnosed with a cerebellar type of MSA. Postmortem delay was ∼30 h. Brain regions with abundant α-Syn inclusions were identified by postmortem histological examination.

For immunohistochemistry (IHC), mouse monoclonal antibody against α-Syn and p62 lck ligand were used. Paraffin sections of human brain tissue were deparaffinated and rehydrated. Pretreatment (cooking in cell conditioning solution 1, pH 8 for 30 min for α-Syn IHC or for 56 min in case of p62 IHC), IHC and counterstaining of nuclei with hematoxylin (Roche) and Bluing reagent (Roche) were performed with the Ventana Bench-Mark XT automated staining system (Ventana) using the UltraView Universal DAB Detection Kit (Roche). For α-Syn, IHC slides were additionally pretreated in 80% formic acid for 15 min after cooking. Slides were coverslipped with Entellan (Merck) mounting medium. Images were recorded with a BX50 microscope (Olympus) using a 40x objective and cellSens software (Olympus).

### Preparation of the sarkosyl-insoluble fraction from MSA patient brain

Preparation of sarkosyl-insoluble fraction was performed as previously described^41^. Briefly, frozen tissue from the basilar part of the pons (1 cm^3^) was homogenized in high salt (HS) buffer (50 mM Tris-HCl pH 7.5, 750 mM NaCl, 10 mM NaF, 5 mM EDTA) with protease and phosphatase inhibitors (Roche) and incubated on ice for 20 min. The homogenate was centrifuged at 100,000 xg for 30 min. The resulting pellet was washed with HS buffer and then re-extracted sequentially with 1 % Triton X-100 in HS buffer, 1 % Triton X-100 in HS buffer and 30 % sucrose, 1 % sarkosyl in HS buffer and finally PBS. The incubation with 1% sarkosyl in HS buffer was performed overnight at 4 °C. The final fraction was sonicated and the presence of α-Syn aggregates was confirmed by immunobloting against phospho S129 α-Syn.

### Cell culture

To create a stable cell line expressing EGFP-α-SynA53T, SH-SY5Y cells were transfected using Lipofectamine 2000 (Thermo Fisher). Cells were cultured in in Dulbecco’s modified Eagle’s medium (DMEM, Biochrom) supplemented with 10 % fetal bovine serum (FBS, GIBCO), 2 mM L-glutamine (GIBCO) and 2,000 μg/ml geneticin for selection. Polyclonal cell lines were generated by fluorescence-activated cell sorting (FACS). Upon selection, cells were cultured in medium supplemented with 200 μg/ml geneticin (Thermo Fisher) and penicillin/streptomycin (Thermo Fisher).

Cells were seeded as described^19^ using 300 nM (monomer) of α-SynA53T PFFs or gold-conjugated WT α-Syn PFFs. In brief, sonicated PFFs were diluted in a mixture of 50 μl of Optimem (Biochrom) and 3 μl of Lipofectamine 2000. Subsequently the suspension was added to 1 ml of cell culture medium.

### Lentivirus packaging

HEK293T cells (632180, Lenti-X 293T cell line, Takara; RRID: CVCL_0063) for lentiviral packaging were expanded to 70-85 % confluency in DMEM Glutamax (+ 4.5 g/L D-Glucose, - Pyruvate) supplemented with 10 % FBS (Sigma), 1 % G418 (Gibco), 1 % NEAA (Thermo Fisher), and 1 % Hepes (Biomol). Only low passage cells were used. For lentiviral production, a three-layered 525cm^2^ flask (Falcon) was seeded and cells were henceforth cultured in medium without G418. On the following day, cells were transfected with the expression plasmid pFhSynW2 (GFP-SynA53T-Flag, Flag-GFP) or FU3a (p62-tagRFP), and the packaging plasmids psPAX2 and pVsVg using TransIT-Lenti transfection reagent (Mirus). The transfection mix was incubated for 20 min at room temperature (RT) and cell medium was exchanged. 10 ml of transfection mix were added to the flask and incubated overnight. The medium was exchanged on the next day. After 48-52 h, culture medium containing the viral particles was collected and centrifuged for 10 min at 1,200 xg. The supernatant was filtered through 0.45 μm pore size filters using 50 ml syringes, and Lenti-X concentrator (Takara) was added. After an overnight incubation at 4 °C, samples were centrifuged at 1,500 xg for 45 min at 4 °C, the supernatant was removed, and the virus pellet was resuspended in 600 μl TBS-5 buffer (50 mM Tris-HCl, pH 7.8, 130 mM NaCl, 10 mM KCl, 5 mM MgCl_2_). After aliquoting, viruses were stored at −80 °C.

### Primary neurons

Primary cortical neurons were prepared from E15.5 CD-1 wild type mouse embryos (breeding line MpiCrlIcr:CD-1). All experiments involving mice were performed in accordance with the relevant guidelines and regulations. Pregnant females were sacrificed by cervical dislocation, the uterus was removed from the abdominal cavity and placed into a 10?cm sterile Petri dish on ice containing dissection medium, consisting of Hanks’ balanced salt solution supplemented with 0.01 M HEPES, 0.01 M MgSO_4_, and 1% penicillin/streptomycin. Embryos of both sexes were chosen randomly. Each embryo was isolated, heads were quickly cut, and brains were removed from the skull and immersed in ice-cold dissection medium. Cortical hemispheres were dissected and meninges were removed under a stereo-microscope. Cortical tissue from typically six to seven embryos was transferred to a 15 ml sterile tube and digested with 0.25 % trypsin containing 1 mM 2,2′,2″,2′′′-(ethane-1,2-diyldinitrilo) tetraacetic acid (EDTA) and 15 μl 0.1 % DNAse I for 20 min at 37 °C. The enzymatic digestion was stopped by removing the supernatant and washing the tissue twice with Neurobasal medium (Invitrogen) containing 5 % FBS. The tissue was resuspended in 2 ml medium and triturated to achieve a single cell suspension. Cells were spun at 130 xg, the supernatant was removed, and the cell pellet was resuspended in Neurobasal medium with 2 % B27 (Invitrogen), 1 % L-glutamine (Invitrogen) and 1 % penicillin/streptomycin (Invitrogen). For immunostaining, neurons were cultured in 24-well plates on 13 mm coverslips coated with 1 mg/ml poly-D-lysine (Sigma) and 1 μg/ml laminin (Thermo Fisher Scientific) (100,000 neurons per well). For MTT assay, neurons were cultured in 96-well plates coated in the same way (19,000 neurons per well). For Cryo-ET, EM grids were placed in 24-well plates and coated as above (120,000 neurons per well). For lentiviral transduction at DIV 10, viruses were thawed and immediately added to freshly prepared neuronal culture medium. Neurons in 24-well plates received 1 μl of virus/well, while neurons in 96 well-plates received 0.15 μl of virus/well. A fifth of the medium from cultured neurons was removed and the equivalent volume of virus-containing medium was added. Three days after transduction, 2 μg/ml of seeds (α-SynA53T PFFs, gold-conjugated WT α-Syn PFFs or MSA-derived aggregates) were added to the neuronal culture medium.

### MTT viability assay

Viability of transduced neurons was determined using Thiazolyl Blue Tetrazolium Bromide (MTT; Sigma-Aldrich). Cell medium was exchanged for 100 μl of fresh medium, followed by addition of 20 μl of 5 mg/ml MTT in PBS and incubation for 2-4 h at 37 °C, 5 % CO_2_. Subsequently, 100 μl solubilizer solution (10 % SDS, 45 % dimethylformamide in water, pH 4.5) was added, and on the following day absorbance was measured at 570 nm. Each condition was measured in triplicates per experiment and absorbance values were averaged for each experiment. Viability values of neurons seeded with α-Syn aggregates were normalized to those of neurons that received PBS only.

### Immunofluorescence

Immunofluorescence stainings on SH-SY5Y cells were performed 24 h after seeding. Cells were fixed for 10 min with 4 % paraformaldehyde (PFA) in PBS and subsequently incubated for 5 min in permeabilization solution (0.1 % Triton X-100 in PBS) at RT. After blocking with 5 % milk in permeabilization solution, primary antibodies were diluted in blocking solution and incubated with the cells over night at 4 °C. Secondary antibodies were incubated with the cells in blocking solution for 3 h at room temperature. The coverslips were subsequently incubated with 500 nM DAPI for 10 min and then mounted on glass slides. Images were taken using a CorrSight microscope (Thermo Fisher) in spinning disc mode with a 63x oil immersion objective.

Primary neurons were fixed with 4% PFA in PBS for 20 min; remaining free groups of PFA were blocked with 50 mM ammonium chloride in PBS for 10 min at RT. Cells were rinsed once with PBS and permeabilized with 0.25 % Triton X-100 in PBS for 5 min. After washing with PBS, blocking solution consisting of 2 % BSA (Roth) and 4 % donkey serum (Jackson ImmunoResearch) in PBS was added for 30 min at RT. Coverslips were transferred to a light protected humid chamber and incubated with primary antibodies diluted in blocking solution for 1 h. Cells were washed with PBS and incubated with secondary antibodies diluted 1:250 in blocking solution, with 0.5 mg/ml DAPI added to stain the nuclei. Coverslips were mounted on Menzer glass slides using Prolong Glass fluorescence mounting medium. Confocal images were obtained at a SP8 confocal microscope (Leica) using 40x or 63x oil immersion objectives (Leica). Neurons containing aggregates in the soma were manually quantified using the Cell Counter plugin of ImageJ^42^ (RRID: SCR_003070).

### Negative stain EM

For negative stain analysis, continuous carbon Quantifoil grids (Cu 200 mesh, QuantifoilMicro Tools) were glow discharged using a plasma cleaner (PDC-3XG, Harrick) for 30 s. Grids were incubated for 1 min with PFFs, blotted and subsequently washed 2 times with water for 10 s. The blotted grids were stained with 0.5 % uranyl acetate solution, dried and imaged in Polara cryo-electron microscope (Thermo Fisher) operated at 300 kV using a pixel size of 2.35 or 3.44 Å.

### Cryo-ET sample preparation

Quantifoil grids (R1/4 or 1.2/20, Au mesh grid with SiO2 film, QuantifoilMicro Tools) were glow discharged using a plasma cleaner (PDC-3XG, Harrick) for 30 s. Cells were plated on the grids as described above. SH-SY5Y cells were seeded with α-Syn aggregates 24 h after plating and plunge frozen after another 24 h. Neurons were transduced on DIV 10, seeded with α-Syn aggregates on DIV 13 and plunge frozen on DIV 20. Plunge freezing was performed on a Vitrobot (Thermo Fisher) with the following settings: temperature, 37 °C; humidity, 80 %; blot force, 10; blot time, 10 s. The grids were blotted from the back and the front using Whatman filter paper and plunged into a liquid ethane/propane mixture. Subsequently the vitrified samples were transferred into cryo-EM boxes and stored in liquid nitrogen.

### Correlative cryo-light microscopy and cryo-FIB milling

Grids were mounted onto autogrid sample carriers (Thermo Fisher) that contain cut-out regions to facilitate shallow-angle FIB milling. Subsequently grids were transferred into the stage of a CorrSight cryo-light microscope (Thermo Fisher) cooled at liquid nitrogen temperature. Overview images of the grids were acquired using a 20x lens (air, N.A., 0.8). Cells containing fluorescence signal of interest (GFP-α-Syn or p62-RFP) were mapped using MAPS software 2.1 (Thermo Fisher; RRID: SCR_018738).

The samples were transferred into a Scios or Quanta dual beam cryo-FIB/scanning electron microscopes (SEM; Thermo Fisher). To avoid charging of the samples, a layer of inorganic platinum was deposited on the grids. That was followed by the deposition of organometallic platinum using an *in situ* gas injection system (working distance - 10 mm, heating - 27 °C, time - 8 s) to avoid damaging the cells by out-of-focus gallium ions. Subsequently 2D-correlation was performed using MAPS and the 3-point alignment method between the fluorescence and the SEM image as described^17^.

For FIB milling, the grid was tilted to 18° and gallium ions at 30 kV were used to remove excess material from above and below the region of interest. Rough milling was conducted at a current of 0.5 nA and fine milling at a current of 50 pA, resulting in 100-200 nm thick lamellas.

### Cryo-ET data collection and reconstruction

The lamellas were transferred into a Titan Krios cryo-electron microscope (Thermo Fisher) operated at 300 kV and subsequently loaded onto a compustage cooled to liquid nitrogen temperatures. Lamellas were oriented perpendicular to the tilt axis. Images were collected using a 4 k x 4 k K2 Summit or K3 (Gatan) direct detector cameras operated in dose fractionation mode (0.2 s, 0.15 e^-^/Å^2^). A BioQuantum (Gatan) post column energy filter was used with a slit width of 20 eV. Tilt series were recorded using SerialEM^43^ (RRID: SCR_017293) at pixel size of 3.52 or 4.39 Å. Tilt series were recorded dose-symmetrically^44^ from −50° to +60° with an angular increment of 2°, resulting in a total dose of 100-130 e^-^/Å^2^ per tilt series. Frames were aligned using Motioncor2^45^. Final tilt series were aligned using fiducial-less patch tracking and tomograms were reconstructed by using back projection in IMOD^46^ (RRID: SCR_003297). Contrast was enhanced by filtering the tomograms using tom_deconv (https://github.com/dtegunov/tom_deconv) within MATLAB (MathWorks).

### Tomogram segmentation

The membranes of the tomograms were segmented using the automatic membrane tracing package TomoSegMemTV^47^. The results were refined manually in Amira (FEI Visualization Science Group; RRID: SCR_014305). The lumen of organelles was filled manually based on the membrane segmentations. For tracing of α-Syn fibrils, the XTracing module^48^ of Amira was used. For that the tomograms were first denoised with a non-local means filter, and subsequently searched with a cylindrical template of 10 nm diameter and 80 nm length. Based on the cross-correlation fields, thresholds producing an optimal balance of true positives and negatives were applied. Filaments were subsequently traced with a search cone of 50 nm length and an angle of 37°. The direction coefficient was 0.3 and the minimum filament length was set to 100 nm. Selected filaments were inspected visually.

### Tomogram analysis

#### Cytosolic fibril density

The density of fibrils within the inclusion was calculated as the fraction of cytosolic volume occupied by fibrils. Cellular volume was calculated multiplying the X and Y dimensions of the tomogram by the thickness of the lamella along the Z direction. To calculate cytoplasmic volume, the lumina of organelles were subtracted from the tomogram volume. Fibril volume was calculated approximating fibrils by cylinders with radius of 5 nm and the length calculated by filament tracing. Calculations were performed in MATLAB.

#### Fibril persistence length

The persistence length (L_p_) measures the stiffness of polymers as the average distance for which a fibril is not bent. It was calculated using an in-house script as previously described^16^ executed in MATLAB. Briefly, L_p_ is calculated as the expectation value of cos θ, where θ is defined as the angle between two tangents to the fibril at positions 0 and l^49^:

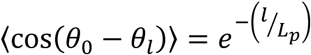

The Young’s modulus (E) can be calculated from L_p_ as:

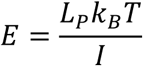

where k_B_ is the Boltzmann constant (1.38 × 10^−23^ m^2^ kg s^-2^ K^-1^), T is the absolute temperature (here 295 K) and I is the momentum of inertia. Approximating the fibril by a solid rod, I can be calculated from its radius r as:

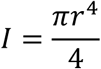

Here we used r = 5 nm.

#### Fibril-membrane distance

Fibril-membrane nearest distance was defined for each point on the fibril as the minimum Euclidean distance to another point on a membrane. The algorithm computing fibril-membrane nearest distances can be summarized as follows:

- For each tomogram:
  - Use the segmentation of organelle lumina to compute the distance transform tomogram^50^, which calculates the Euclidean distance from each background voxel to the nearest segmented one.
  - For each fibril:
    - The curve defined by Amira’s Xtracing module during segmentation is sampled uniformly each 5 nm (i.e. similar to the fibril radius).
    - For each point in the fibril:
      - To achieve subvoxel precision, get the interpolated value of the distance transform tomogram at the coordinates of that point.
      - Add this value to a list of fibril-membrane nearest distances.

The probability density was computed as the normalized histogram of the list of fibril-membrane nearest distances.

To test whether these fibril-membrane nearest distances resulted from random or specific interactions, we compared the experimentally determined distances with those of simulated fibrils. These simulated fibrils were created by randomly shifting and rotating the experimentally measured fibrils as follows:

- For each tomogram:
  - Generate 200 synthetic tomograms:
    - Take randomly an input experimental fibril as reference.
    - Shift the reference fiber in respect to its center at a random distance in a range of [10, 20] nm.
    - Rotate the fibril randomly with respect to the fibril center with Euler angles selected randomly in the range of [0, 10] degrees.
    - Try to insert the resulting fibril in the synthetic tomogram. The insertion fails in the following cases:
      - The fibril intersects with another one, considering that fibrils have a cross-section radius of 5 nm.
      - The fibril intersects with a segmented membrane.
      - Part of the fibril is out of the tomogram boundaries.
    - Iterate until 50 fibrils are inserted or 5000 tries are reached.

#### Inter-membrane distance

Each segmented lumen was labeled differently to identify different organelles. The inter-membrane nearest distance for a point on a membrane was defined as the minimum Euclidean distance to another point on a membrane associated to a different lumen. The algorithm for computing inter-membrane nearest distances can be summarized as follows:

- For each tomogram:
  - Assign labels for the lumen of each organelle.
  - Associate segmented membranes and lumina by a proximity criterion. For each voxel in a membrane segmentation, the label of the nearest lumen voxel is determined. The lumen is then associated to the membrane segmentation most frequently found.
  - For each lumen:
    - Compute the distance transform tomogram^50^ from all lumina.
    - Erase the current lumen.
    - For each pixel on the membrane segmentation associated to the current lumen:
      - Get the interpolated value of the distance transform tomogram at the coordinates of that point.
      - Add this value to a list of inter-membrane nearest distances.

Probability densities were computed as described for fibril-membrane nearest distances.

### Statistical analysis

For the quantification of the percentage of neurons with aggregates using light microscopy (Extended Data Fig. 1e), N = 4 (GFP-α-Syn + PFFs) and 3 (endogenous α-Syn + PFFs) independent experiments were performed, and a total of 100-500 neurons per condition and per experiment were counted. Statistical analysis was carried out by two-tailed unpaired t-test with Welch’s correction in Prism 6 (GraphPad; RRID: SCR_002798).

For the quantification of neuronal viability using the MTT assay (Extended Data Fig. 1f), N = 3 independent experiments were performed for all conditions. Untransduced and unseeded control cells were used as reference. Statistical analysis was carried out by two-way ANOVA and Dunnett’s multiple comparison test in Prism 6.

The number of tomograms and fibrils, as well as the total membrane area analyzed for each condition are shown in Extended Data Table 1.

Statistical analysis of cytosolic fibril density (Fig. 2e) was carried out by one-way ANOVA. Confidence intervals for fibril-membrane (Fig. 4b) and inter-membrane (Fig. 4b) distances were calculated as the 5-95 percentiles from the curves of each individual tomogram. The differences between the curves within 20 nm were statistically analyzed by Kolmogorov-Smirnov test.

Additional information on statistical analyses can be found in the Source Data files.

## Data and code availability

All data supporting the findings of this study are available within this paper. Source Data for Fig. 2d, e, Fig. 4b, c, Extended Data Fig. 1e, f, Extended Data Fig. 4, FACS (Supplementary Fig. 1) and gel source images (Supplementary Fig. 2) are available with the online version of this paper. The tomograms shown in Fig. 1 and Fig. 2 are available in EMPIAR through accession codes EMD-11401 (Fig. 1a), EMD-11417 (Fig. 1e) and EMD-11416 (Fig. 2a).

The tomogram deconvolution filter is available at: https://github.com/dtegunov/tom_deconv.

The script for the calculation of L_p_ is available at: https://github.com/FJBauerlein/Huntington.

The scripts for fibril-membrane and inter-membrane distance calculations were performed within the PySeg software^51^ and are available at: https://github.com/anmartinezs/pyseg_system/tree/master/code/pyorg/scripts/filaments.

## Acknowledgments

We thank Philipp Erdmann, Günter Pfeifer, Jürgen Plitzko and Miroslava Schaffer for electron microscopy support, Ana Jungclaus and Nadine Wischnewski for wet lab support. We thank Dieter Edbauer, Hilal Lashuel, David Rubinsztein and Didier Trono for sharing plasmids. We thank Konstanze Winklhofer and Joerg Tazelt for sharing the SH-SY5Y cell line and helpful discussions. We are also grateful to Sophie Keeling for help in plasmid cloning, Javier Collado for help with sample preparation, Jonathan Schneider for help with image processing, Patrick Auer for help in aggregate quantification, as well as Itika Saha, Eri Sakata and Patricia Yuste-Checa for helpful discussions. Finally, we thank the anonymous MSA patient and his family for the donation of brain tissue to the Neurobiobank Munich. Fluorescence-activated cell sorting was carried out at the Imaging Facility of the Max Planck Institute of Biochemistry. V.A.T. was supported by the Graduate School of Quantitative Biosciences Munich. V.A.T., I.R.-T., A.M.-S., F.J.B., Q.G., W.B., I.D., M.S.H., F.U.H. and R.F.-B. have received funding from the European Commission (FP7 GA ERC-2012-SyG_318987-ToPAG). I.D. acknowledges financial support from the Horst Kübler-Stiftung. V.A.T., T.A., M.S.H., F.U.H. and R.F.-B. acknowledge funding from the Deutsche Forschungsgemeinschaft (DFG, German Research Foundation) through Germany’s Excellence Strategy - EXC 2067/1-390729940 (R.F.-B.) and EXC 2145 – 390857198 (V.A.T., T.A., M.S.H. and F.U.H).

## Author contributions

V.A.T. performed biochemical and electron microscopy experiments, immunofluorescence imaging of SH-SY5Y cells and contributed to computational data analysis. I.R.-T. produced lentivirus and neuronal cultures, and performed viability assays and immunofluorescence imaging of neurons. A.M.S. developed software procedures for data analysis. F.B. and Q.G. contributed to data analysis. T.A. collected the autopsy case, characterized it neuropathologically and performed immunohistochemistry. V.A.T., I.R.-T., W.B., I.D., M.S.H., F.U.H. and R.F.-B. planned research. I.D. supervised neuronal culture experiments. M.S.H. and F.U.H. supervised biochemical experiments. R.F.-B. supervised electron microscopy experiments and data analysis. R.F.-B. wrote the manuscript with contributions from all authors.

## Additional information

Supplementary Information is available for this paper. Correspondence and requests for materials should be addressed to F.U.H. and R.F.-B.

## Extended data

**Extended Data Table 1.** Statistics of cryo-ET experiments on mouse neurons. Neurons were either transduced with GFP-α-syn and seeded with PFFs (“GFP-α-syn + PFFs”), transduced with p62-RFP and seeded with PFFs (“Endogenous α-syn + PFFs”), transduced with GFP-α-syn and seeded with aggregates derived from an MSA patient brain (“GFP-α-syn + MSA”), or untransduced and unseeded as control (“Untransduced - PFFs”). The column “Experiments” lists biologically independent replicates. “Analyzed filaments” includes all filaments analyzed in Fig. 2d, e, Fig. 4 and Extended Data Fig. **7**d. “Analyzed membrane area” includes all membranes analyzed in Fig. 4 and Extended Data Fig. **7**e.

| Condition | Experiments | Analyzed tomograms | Analyzed filaments | Analyzed membrane area ($\mu\text{m}^2$) |
| --- | --- | --- | --- | --- |
| GFP-α-syn + PFFs | 3 | 5 | 1471 | 3.83 |
| Endogenous α-syn + PFFs | 3 | 4 | 220 | 2.75 |
| GFP-α-syn + MSA | 3 | 5 | 721 | 3.70 |
| Untransduced - PFFs | 2 | 5 | - | 3.69 |

**Extended Data Fig. 1.**
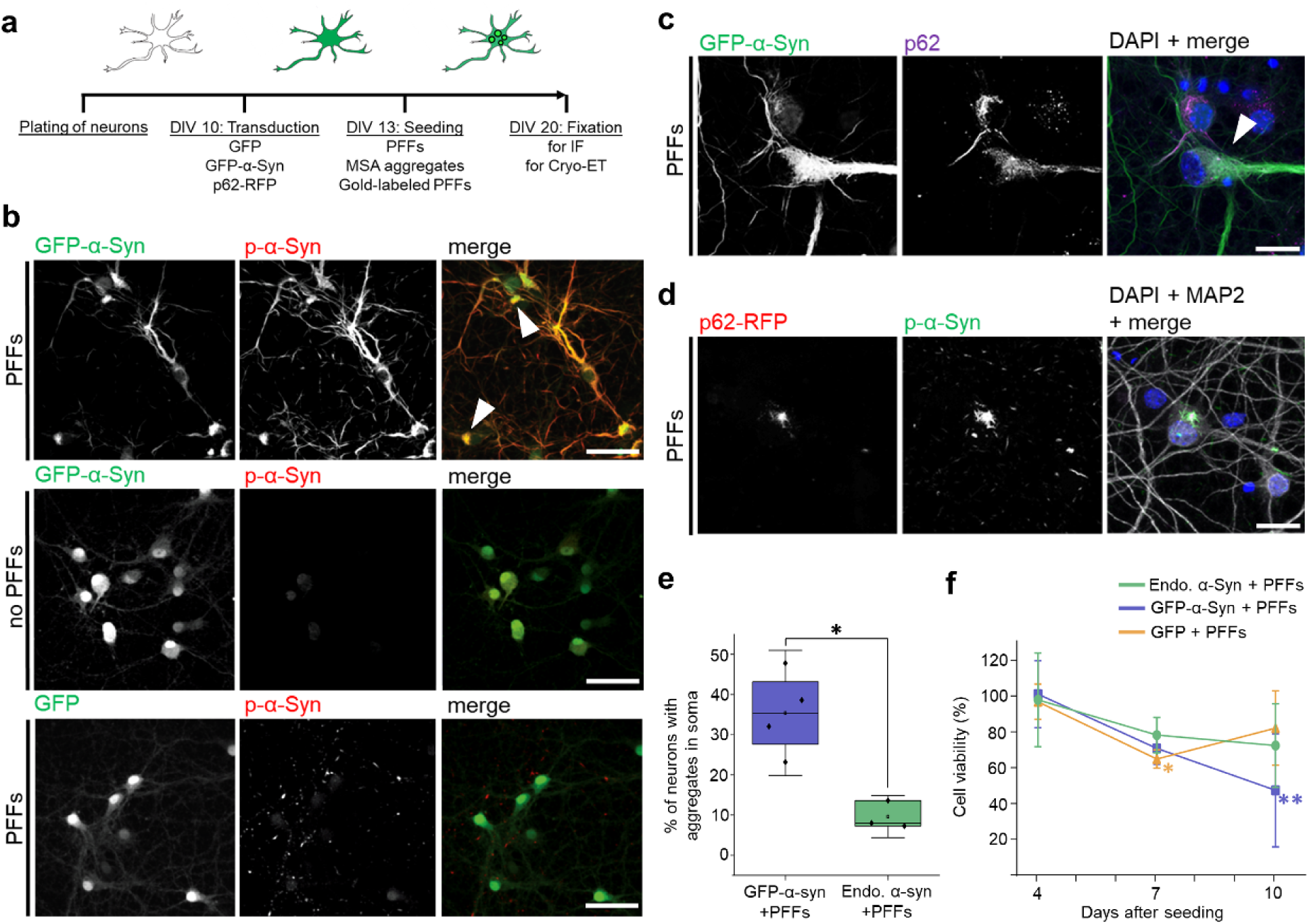
Seeding of α-Syn aggregates in neurons. **a**, Schematic of the seeding of α-Syn aggregates in primary neurons. Primary mouse neurons were cultivated and transduced at day *in vitro* (DIV) 10 with GFP, GFP-α-Syn or p62-RFP. Seeds (PFFs or MSA brain-derived) were applied at DIV 13, and α-Syn inclusions were studied at DIV 20 by light microscopy or cryo-ET upon chemical or cryo-fixation, respectively. For light microscopy imaging, GFP signal was enhanced by staining with an antibody against GFP. **b**, Immunofluorescence imaging of α-Syn aggregates, as detected by an antibody against phosphorylated α-Syn Ser129 (p-α-Syn). Top: aggregate formation (arrowheads) upon seeding cells expressing GFP-α-Syn with exogenous PFFs. Middle: no aggregate formation in cells expressing GFP-α-Syn in the absence of PFFs. Bottom: PFFs seed smaller aggregates in cells with endogenous α-Syn levels that express GFP only as control (see Extended Data Fig. 1e for quantification). Scale bars: 50 μm. **c**, Immunofluorescence imaging of GFP-α-Syn aggregates detected by an antibody against p62. The merged image shows a superposition of the GFP-α-Syn (green), p62 (magenta) and DAPI (blue) channels. An arrowhead indicates the colocalization of GFP-α-Syn and p62. Scale bar: 20 μm. **d**, Immunofluorescence imaging of endogenous α-Syn aggregates positive for p-α-Syn colocalizing with p62-RFP. The merged image shows a superposition of the p62-RFP (red), phospho-α-Syn (green), the neuronal marker MAP2 (grey) and DAPI (blue) channels. Scale bar: 20 μm. **e**, Quantification of the percentage of neurons with aggregates in the soma upon treatment with PFFs of cells transduced with GFP-α-Syn (blue) or untransduced (green; endogenous α-Syn). The horizontal lines of each box represent 75% (top), 50% (middle) and 25% (bottom) of the values, and a black square the average value. Whiskers represent standard deviation and black diamonds the individual data points. * indicates p = 0.011 by two-tailed unpaired t-test with Welch’s correction, N = 4 (GFP-α-Syn + PFFs) and 3 (endogenous α-Syn + PFFs) independent experiments. **f**, Quantification of neuronal viability upon seeding with PFFs for cells expressing endogenous α-Syn (Endo. α-Syn + PFFs), or transduced with GFP-α-Syn (GFP-α-Syn + PFFs) or with GFP only (GFP + PFFs) relative to untransduced and unseeded control cells. Points represent average values and the error bars the standard deviation. *and ** respectively indicate p = 0.04 and p = 0.002 by two-way ANOVA and Dunnett’s multiple comparison test, N = 3 independent experiments for all conditions.

**Extended Data Fig. 2.**
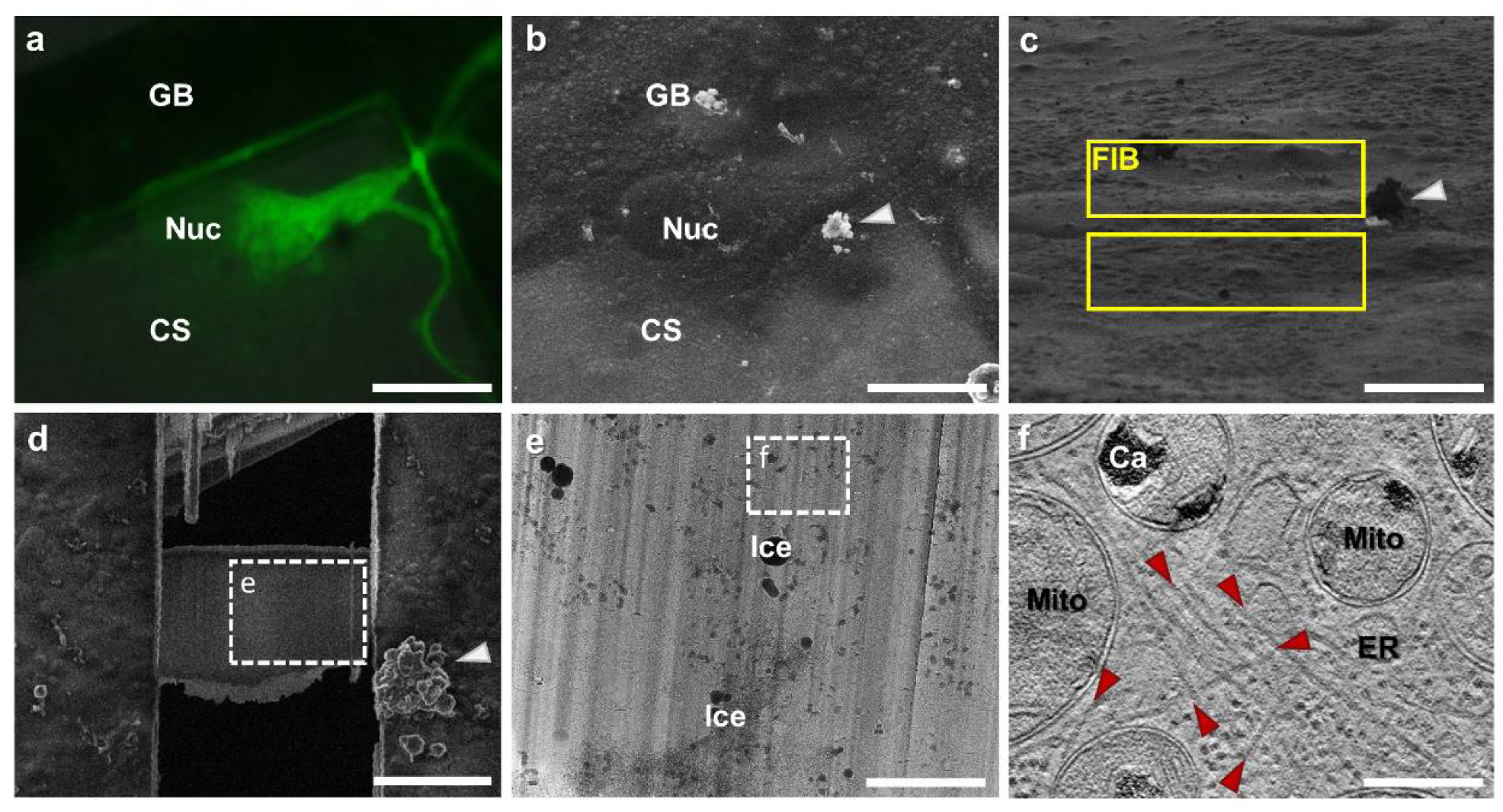
Cryo-ET workflow. **a**, Cryo-light microscopy imaging of GFP fluorescence in a primary neuron grown on the carbon support (CS) of an EM grid. The cell was transduced with GFP-α-Syn at DIV 10 and aggregate formation was seeded by PFFs at DIV 13. The grid was vitrified at DIV 20. GB: grid bar, Nuc: nucleus. Scale bar: 35 μm. **b**, Correlative scanning electron microscopy imaging of the same cell within the cryo-FIB instrument upon coordinate transformation. A white arrowhead marks a piece of ice crystal contamination that can also be found in panels **c** and **d** as visual reference. Scale bar: 35 μm. **c**, FIB-induced secondary electron image of the same cell. Yellow boxes indicate the regions to be milled away by the FIB during lamella preparation. Scale bar: 20 μm. **d**, Scanning electron microscopy imaging of the same cell upon preparation of a 150 nm-thick electron transparent lamella. The white square marks the region of the lamella shown in **e**. Scale bar: 15 μm. **e**, Low magnification transmission electron microscopy image of the area of the lamella marked in **d**. Ice: ice crystal contamination on the lamella surface. The white square marks the region shown in **f**. Scale bar: 3 μm. **f**, A tomographic slice (thickness 1.4 nm) recorded in the area indicated in **e**. Ca: mitochondrial calcium stores, ER: endoplasmic reticulum, Mito: mitochondrion. Red arrowheads indicate α-Syn fibrils. Scale bar: 300 nm.

**Extended Data Fig. 3.**
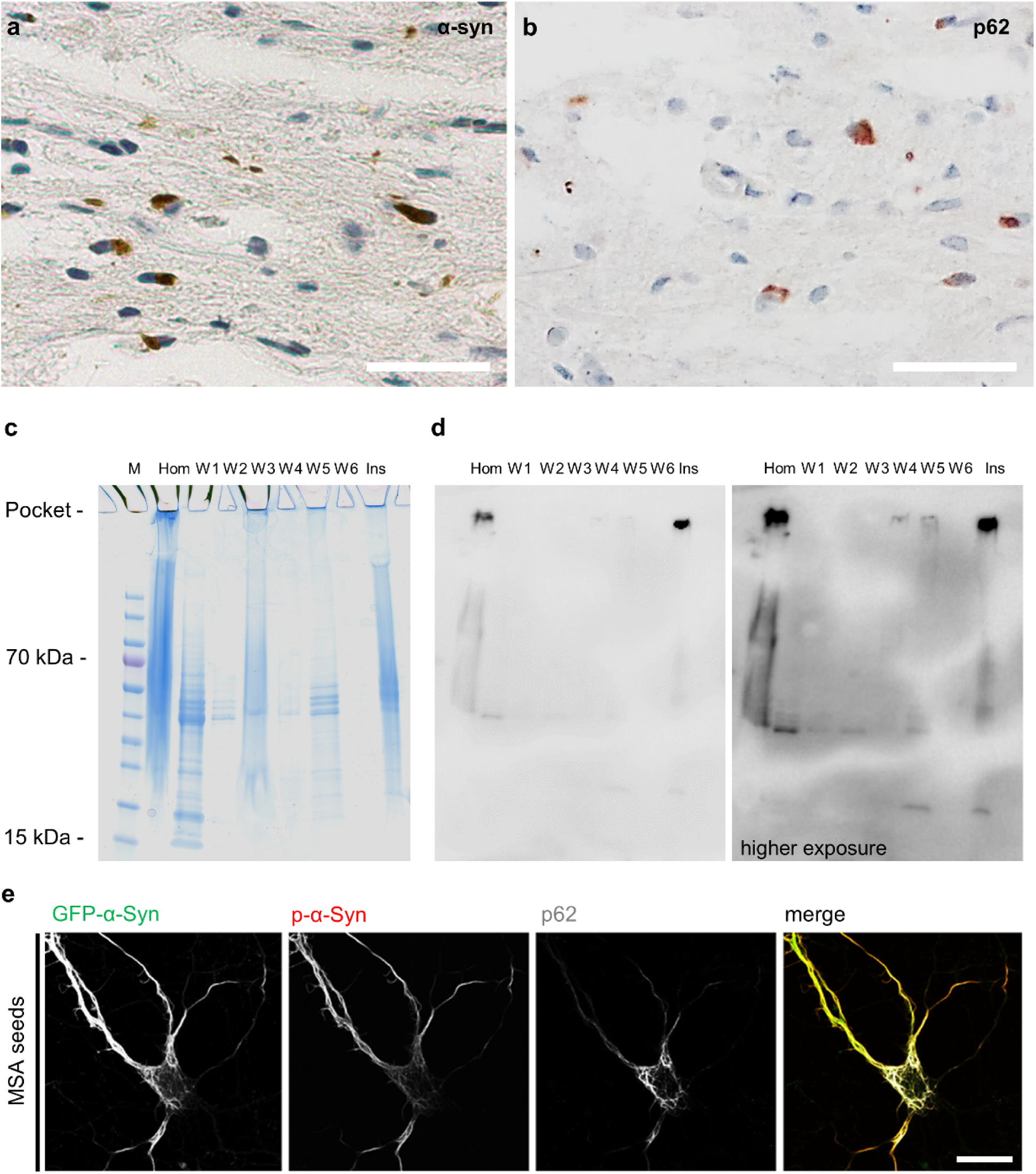
Purification of α-Syn aggregates from MSA patient brain. **a, b**, Immunohistochemistry staining showing cytoplasmic inclusions (brown) positive for α-Syn (**a**) and p62 (**b**) in the basilar part of the pons of the brain of an MSA patient. Aggregates for seeding neurons for cryo-ET imaging were purified from the same region (**c, d**). Scale bars: 50 μm. **c, d**, Purification of α-Syn aggregates from the MSA patient brain shown in **a, b**. Coomassie staining (**c**) and anti-phospho-α-Syn western blot (**d**) of SDS PAGE gels loaded with brain homogenate (Hom), washing fractions (W1-6) and the final sarkosyl-insoluble fraction (Ins) at low (left) and high (right) exposure levels. M: molecular weight marker. Note the aggregated material in the stacking gel. For gel source images, see Supplementary Fig. 2. **e**, Immunofluorescence images of a GFP-α-Syn-expressing neuron seeded with the sarkosyl-insoluble fraction from MSA patient brain, showing aggregates positive for phospho-α-Syn and p62. GFP signal was enhanced by staining with an antibody against GFP. The merged image shows a superposition of the GFP-α-Syn (green), phospho-α-Syn (red) and p62 (grey) channels. Scale bar: 20 μm.

**Extended Data Fig. 4.**
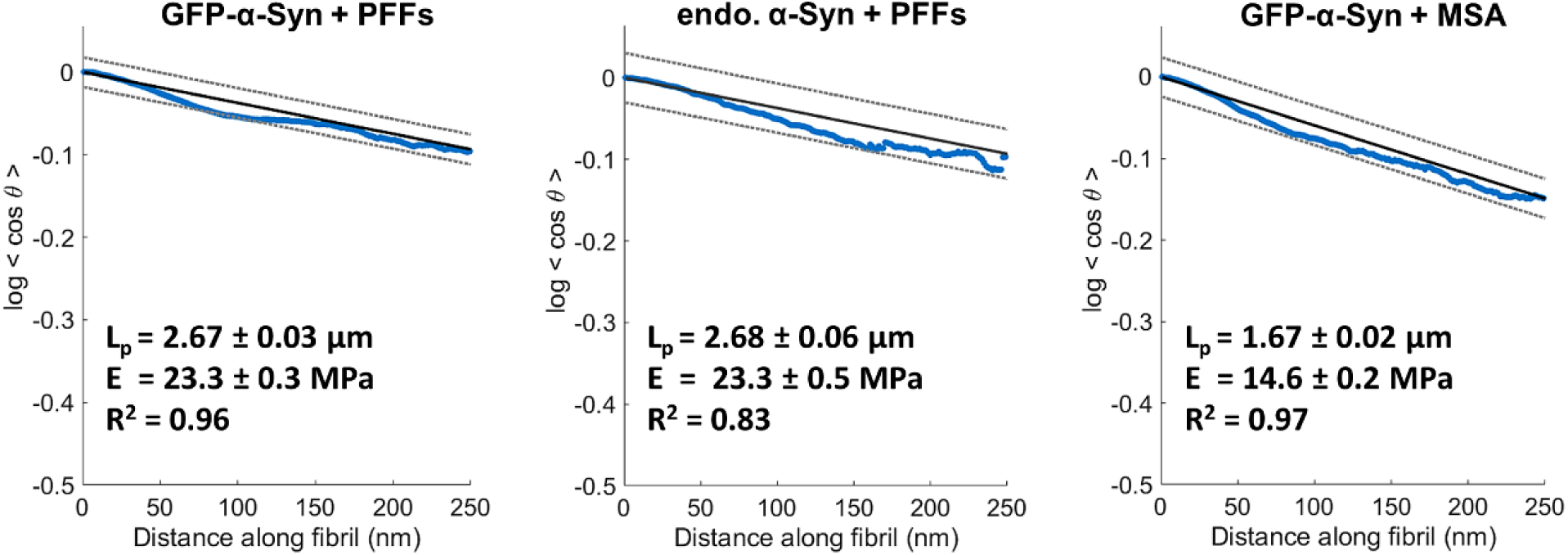
Persistence length of α-Syn fibrils. Linear fit of the total persistence length for all fibrils analyzed (N = 1295 (GFP-α-Syn + PFFs), 220 (endogenous α-Syn + PFFs) and 721 (GFP-α-Syn + MSA) fibrils in total). The blue curves represent the original data. 95% confidence interval (dotted lines) and the values of the persistence length (L_p_), Young’s modulus (E) and coefficients of determination (R^2^) are indicated. Note that the values are almost identical for GFP-α-Syn and endogenous α-Syn seeded with PFFs, but lower for GFP-α-Syn seeded with MSA patient aggregates.

**Extended Data Fig. 5.**
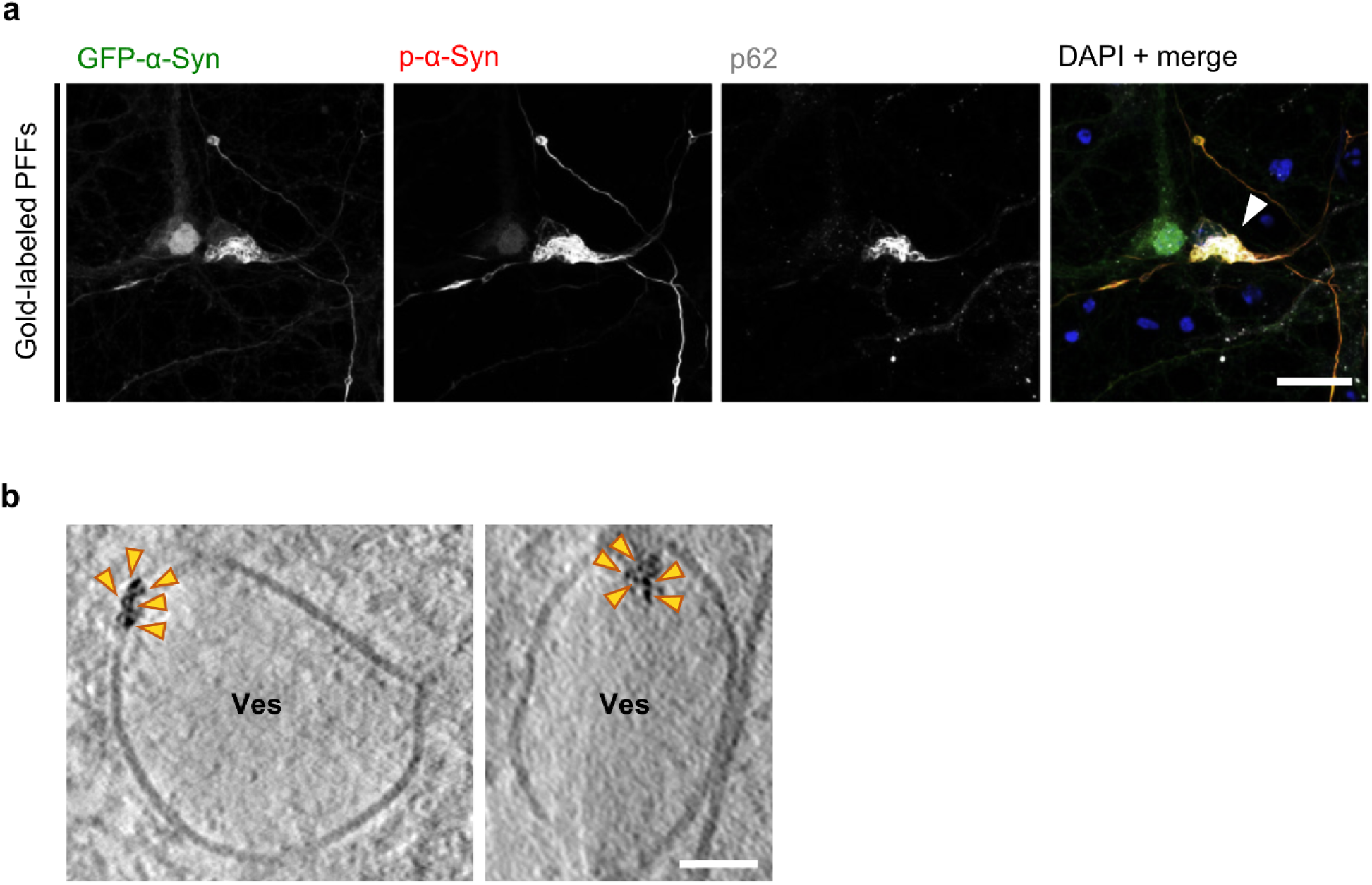
Seeding of α-Syn aggregates in neurons by gold-labeled PFFs. **a**, Immunofluorescence images of a GFP-α-Syn-expressing neuron seeded with gold-labeled PFFs. The cells develop α-Syn aggregates, as detected by antibodies against phosphorylated α-Syn Ser129 and p62. GFP signal was enhanced by staining with an antibody against GFP. The merged image shows a superposition of the GFP-α-Syn (green), phospho-α-Syn (red), p62 (grey) and DAPI (blue) channels. An arrowhead indicates the GFP-α-Syn aggregates. Scale bar: 20 μm. **b**, Tomographic slices (thickness 1.4 nm) showing accumulations of gold particles (orange arrowheads) at the membrane (left) or in the lumen (right) of intracellular vesicles. Ves: vesicles. Scale bar: 50 nm.

**Extended Data Fig. 6.**
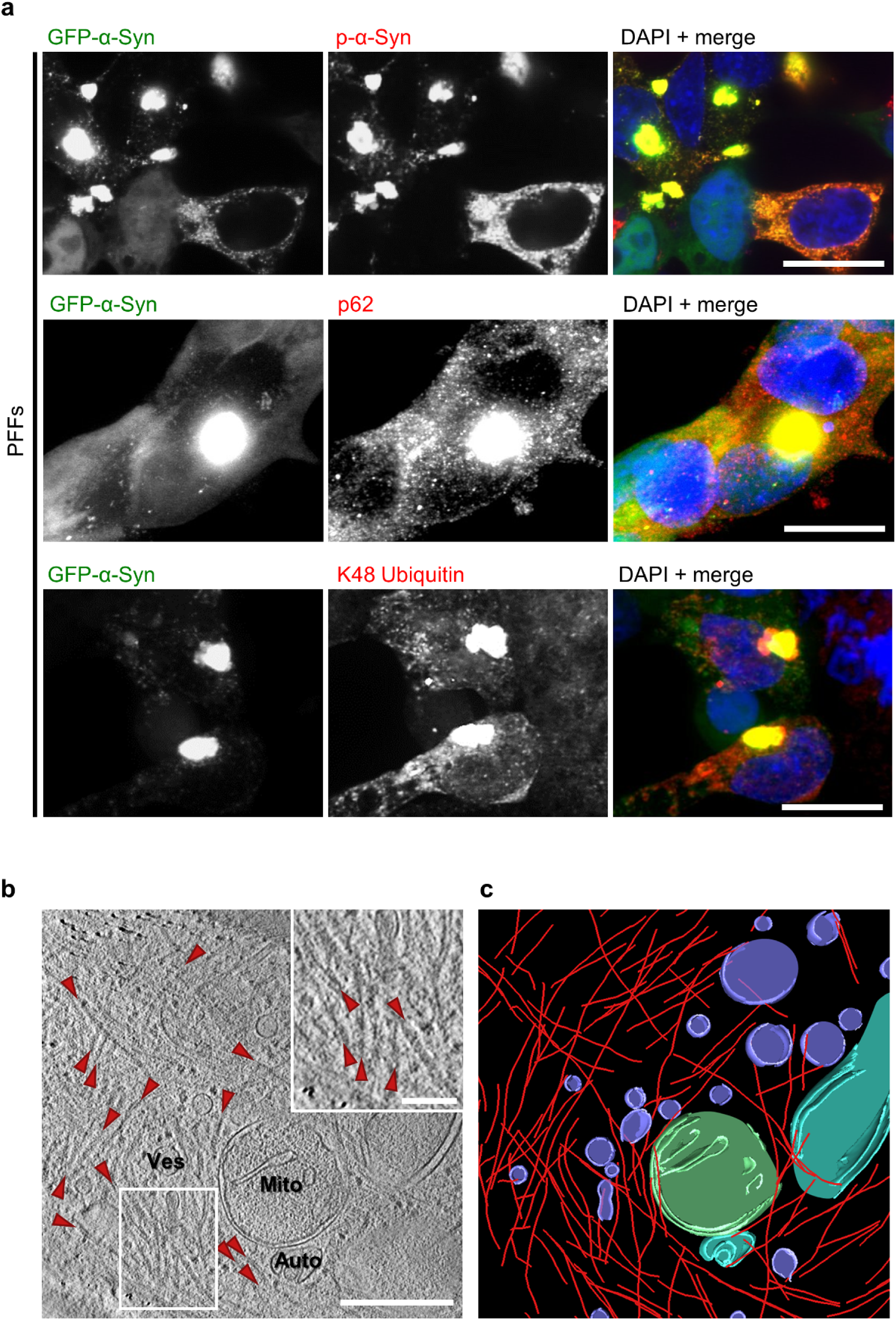
α-Syn aggregates in SH-SY5Y cells. **a**, Immunofluorescence images of SH-SY5Y cells stably expressing GFP-α-Syn and seeded with PFFs. The cells develop α-Syn inclusions, as detected by antibodies against phosphorylated α-Syn Ser129 (top), p62 (middle) and K48-linked ubiquitin (bottom). The merged images show a superposition of the respective green and red channels plus DAPI (blue). Scale bars: 20 μm. **b**, A tomographic slice (thickness 1.8 nm) of an inclusion seeded by PFFs in a SH-SY5Y cell expressing GFP-α-Syn. Auto: autophagosome; Mito: mitochondrion; Ves: vesicles. Fibrils are marked by red arrowheads. Scale bars: 350 nm (main panel) and 100 nm (inset). **c**, 3D rendering of **b** showing α-Syn fibrils (red), autophagosomes (cyan), mitochondria (green) and various vesicles (purple).

**Extended Data Fig. 7.**
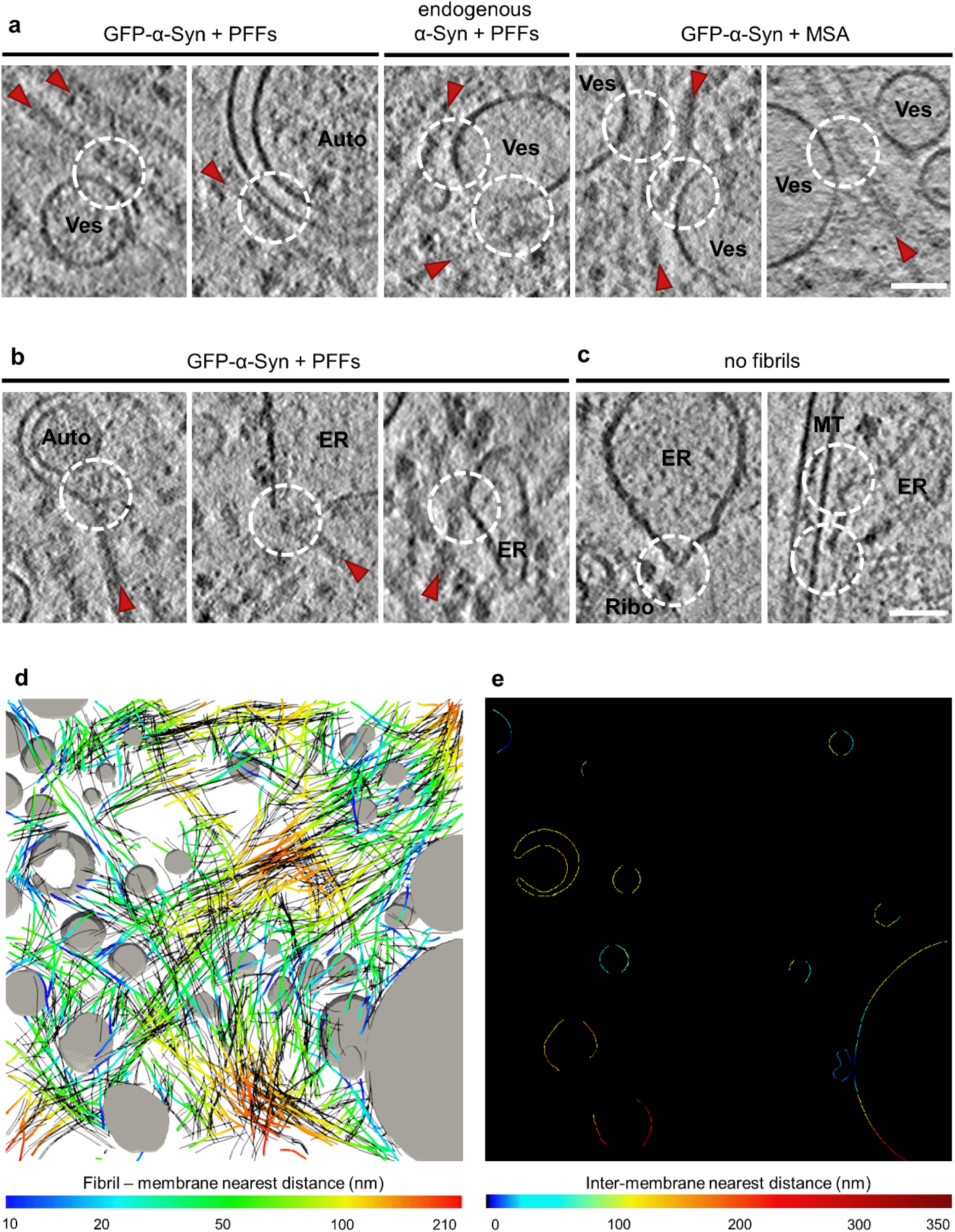
Proximity of α-Syn fibrils and cellular membranes. **a**, Gallery of tomographic slices showing close proximity events (dashed white circles) between α-Syn fibrils (red arrowheads) and different cellular membranes with no apparent interactions. Auto: autophagosome, Ves: vesicles. Tomographic slices are 1.8 nm (GFP-α-Syn + PFFs) or 1.4 nm (endogenous α-Syn + PFFs and GFP-α-Syn + MSA) thick. Scale bar: 60 nm. **b**, Gallery of tomographic slices (thickness 1.8 nm) showing apparent contacts between α-Syn fibrils and different cellular membranes at sites of high membrane curvature (dashed white circles). ER: endoplasmic reticulum. Scale bar: 60 nm. **c**, Tomographic slices showing sites of high membrane curvature (dashed white circles) in the absence of α-Syn fibrils. MT: microtubule; Ribo: ribosome. Tomographic slices are 1.8 nm (left, GFP-α-Syn + PFFs) or 1.4 nm (right, endogenous α-Syn + PFFs) thick. Scale bar: 60 nm. **d**, 3D rendering shown in Fig. 1d and Fig. 4a with α-Syn fibrils color-coded according to their distance to the nearest cellular membrane (grey). To elucidate whether the events of close proximity between fibrils and membranes were caused by chance or mediated by molecular interactions, random shifts (by 10 – 20 nm) and rotations (between 0 and 10°) were performed to the experimentally determined location of the fibrils. Black lines show 5 simulations for 50 randomly chosen fibrils. **e**, Measurements of inter-membrane distances for a 2D slice of the tomogram shown in **d**.

